# Regulatory T cells control the dynamic and site-specific polarization of total CD4 T cells following *Salmonella* infection

**DOI:** 10.1101/710665

**Authors:** Slater L. Clay, Alberto Bravo-Blas, Daniel M. Wall, Megan K.L. MacLeod, Simon W. F. Milling

**Affiliations:** Institute of Infection, Immunity and Inflammation, College of Medical, Veterinary and Life Sciences, University of Glasgow, 120 University Place, Glasgow G12 8TA, United Kingdom; Department of Immunology and Infectious Diseases, Harvard T.H. Chan School of Public Health, 665 Huntington Avenue, Boston, MA 02115, USA; Institute of Cancer Sciences, University of Glasgow and Cancer Research UK Beatson Institute, Glasgow, G61 1BD, United Kingdom

## Abstract

FoxP3^+^ regulatory T cells (Tregs) control inflammation and maintain mucosal homeostasis, but their functions during infection are poorly understood. Th1, Th2 and Th17 cells can be identified by master transcription factors (TFs) T-bet, GATA3 and RORγT; Tregs also express these TFs. While T-bet^+^ Tregs can selectively suppress Th1 cells, it is unclear whether distinct Treg populations can alter Th bias. To address this, we used *Salmonella enterica* serotype Typhimurium to induce non-lethal colitis. Following infection, we observed an early colonic Th17 response within total CD4 T cells, followed by a Th1 bias. The early Th17 response, which contains both Salmonella-specific and non-*Salmonella*-specific cells, parallels an increase in T-bet^+^ Tregs. Later, Th1 cells and RORγT^+^ Tregs dominate. This reciprocal dynamic may indicate that Tregs selectively suppress Th cells, shaping the immune response. Treg depletion 1-2 days post-infection shifted the early Th17 response to a Th1 bias; however, depletion 6-7 days post-infection abrogated the Th1 bias. Thus, Tregs are necessary for the early Th17 response, and for a maximal Th1 response later. These data show that Tregs shape the overall tissue CD4 T cell response and highlight the potential for subpopulations of Tregs to be used in targeted therapeutic approaches.

## INTRODUCTION

*Salmonella enterica* serovar Typhimurium (*S.* Tm) infects a wide range of animals and is a common non-typhoidal *Salmonella* serotype in humans. To study intestinal *S.* Tm infection, a streptomycin pre-treatment model has been developed, creating a niche for *S.* Tm to colonize^1,2^. This facilitates consistent intestinal infection and colitis^3,4^. Because the use of virulent *S.* Tm strains in susceptible mice causes high mortality, we here combine streptomycin pre-treatment with an attenuated *S.* Tm strain that induces consistent non-lethal colitis. This enabled analysis of the CD4 T cell response in the colonic lamina propria and the draining lymph nodes.

CD4 T helper (Th) cells drive intestinal immune responses and can express master transcription factors (TFs). T-bet, GATA3 and RORγT are used to identify Th1, Th2 and Th17 cells, respectively. CD4 T cells are critical for both clearance of *S.* Tm infection and protection from re-infection^5,6^. In particular, Th1 cells are required for *S.* Tm clearance and protective immunity in systemic infection^5,7–9^. In addition to Th1 cells, an early Th17 response has been identified in the intestines, which is important for limiting epithelial damage and systemic dissemination^10–12^. Thus, an effective immune response to *S.* Tm involves both Th17 and Th1 cells. It is unclear however, how the balance between these responses is controlled.

FoxP3^+^ regulatory T cells (Tregs) regulate conventional T cells (Tconvs) and control the CD4 T cell response during infection^13–15^. Tregs can differentiate in the thymus (tTregs) or peripheral tissue (pTregs). While both tTregs and pTregs are found in the intestines^16–18^, their contribution to immune responses to infection is poorly characterized. In addition to FoxP3, Tregs can express Th master TFs T-bet, RORγT or GATA3^19–22^. The role of these Treg TFs is the subject of ongoing research and accumulating evidence suggests that Tregs expressing Th TFs may selectively suppress their respective Th subset^21,23,24^. T-bet^+^ Tregs can selectively target Th1 cells, but it is unclear whether Tregs expressing other TFs have similarly selective functions^21,24^. It is also unclear whether selective suppression can alter the balance of Th cell responses during infection.

To investigate the role of Tregs in controlling the CD4 T cell response, we employed a model of *S.* Tm colitis with an attenuated strain allowing identification of an *S*. Tm-specific T cell clonotype. Characterization of Tconvs and Tregs highlighted a dynamic colonic T cell response within total T cells at the infection site. This is dominated initially by Th17 cells, followed by a sustained Th1 response. Interestingly, we identified a reciprocal dynamic between Tconvs and Tregs expressing the same TFs. This reciprocity indicated that subpopulations of Tregs may selectively suppress Th subsets. Treg depletion experiments were therefore carried out using FoxP3^DTR^ mice^25^. Results show that Tregs are necessary for the early Th17 response and contribute to the later Th1 bias. Thus, Tregs not only inhibit CD4 T cells, but shape the T cell response in a fine-tuned, tissue-specific manner.

## RESULTS

### *Salmonella* infection with BRD509-2W1S causes non-lethal colitis with transient weight loss and increased numbers of total and 2W1S-specific CD4 T cells

To investigate the CD4 T cell response to *S.* Tm in the intestines, attenuated strain BRD509-2W1S was orally administered to C57BL/6 mice 24 hrs following streptomycin treatment. Infection caused non-lethal colitis and allowed identification of *S.* Tm-specific T cells. Representative images of large intestines, including the caecum and colon, are shown from infected (top) and uninfected (bottom) animals 6 days post-infection (p.i.) (Fig. 1a). Infected tissues show colonic shortening, edema, empty and shriveled caeca and an enlarged caecal patch (Fig. 1a, inset). Despite physical signs of colitis, *S.* Tm-infected mice lost <5% of initial weight, which recovered to levels of mock (PBS)-infected controls by 14 days p.i. (Fig. 1b). To assess bacterial dissemination, intestinal and lymphoid tissues were processed and plated. At day 6 p.i. *S.* Tm CFUs were recovered from the colon, mesenteric lymph nodes (MLN), Peyer’s patches (PP) and small intestine (SI); significantly more CFUs were recovered from the colon and MLN than the SI (Fig. 1c).

**Fig. 1.**
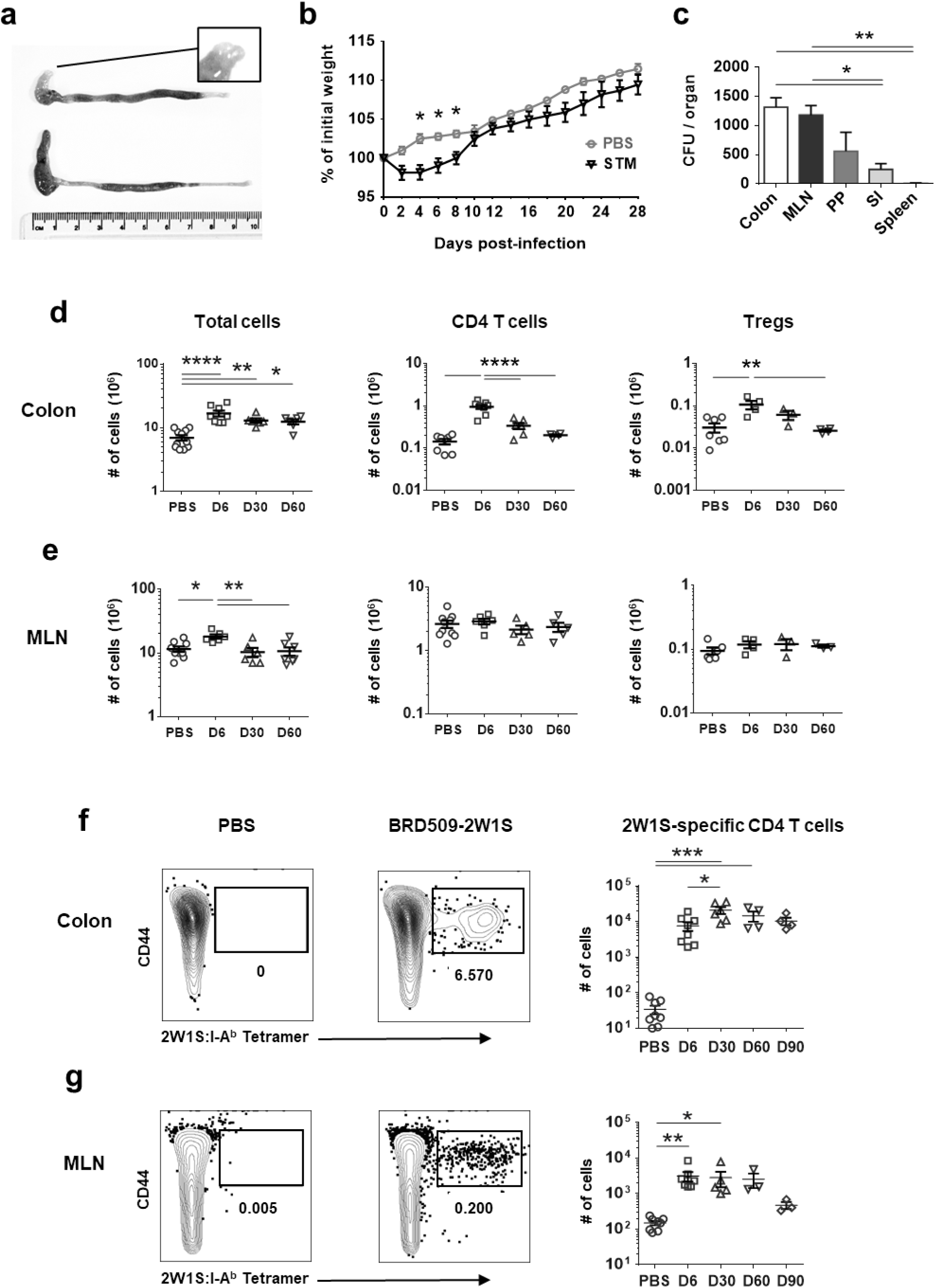
Oral administration of *S.* Tm strain BRD509-2W1S induces non-lethal intestinal infection, colitis and *S.* Tm-specific CD4 T cells in the colon and MLN. **a** Representative images of colon and caeca from infected (top) and uninfected (bottom) mice, 6 days p.i. Inset shows the tip of infected caecum with enlarged caecal patch. **b** Change in initial weight is plotted following infection with BRD509-2W1S or mock infection with PBS. **c** *S.* Tm CFUs recovered from five tissues 6 days p.i. are plotted. **d** The number of total colonic cells, CD4 T cells and Tregs are plotted from PBS (mock)-infected and *S.* Tm-infected mice at day 6, 30 and 60 p.i. Total cells are plotted as enumerated from single cell suspension by hemocytometer. Colonic CD4 T cells were identified by flow cytometry as live, single CD45^+^, CD3^+^, Dump^**-**^ (MHCII, B220, CD8, CD64) CD4^+^ cells and Tregs were identified as FoxP3^+^ CD4 T cells. **e** Total MLN cells, CD4 T cells and Tregs were identified and plotted at the same timepoints as above. **f** Representative plots show the proportion of CD4 T cells that are labelled with 2WIS:1-A^b^ tetramers in mock-infected (left) and BRD509-2W1S-infected samples (middle). The number of 2W1S-specific colonic CD4 T cells are plotted (right) at day 6, 30, 60 and 90 p.i. **g** Representative plots and charts of 2W1S-specific CD4 T cells in the MLN are displayed as above. Weight charts (**b**) show data from two independent experiments (*n*=3-5) and CFU data (**c**) is from one experiment (*n*=3). Data points (**d-g**) represent individual animals from 2 independent experiments (*n*=3-11). In all charts, mean ± standard error of the mean (SEM) are plotted. Statistical significance between weights at specific time points are calculated by multiple Student’s t tests. Statistical difference between all other groups are calculated by one-way ANOVA with Tukey’s test. ns, not significant; \**p*<.05; \*\**p*<.01; \*\*\**p*<.001; \*\*\*\**p*<.0001.

Next, changes in total cell numbers, CD4 T cells and FoxP3^+^ Tregs were enumerated in the colonic lamina propria (LP) (Fig. 1d) and MLN (Fig. 1e) at days 6, 30 and 60 p.i. There was a significant increase in the number of total cells, CD4 T cells and Tregs at day 6 p.i., and total cell numbers remained elevated until 60 days p.i. (Fig. 1d). The number of total MLN cells also increased 6 days p.i., but this increase was resolved by day 30 p.i. Unlike in the colon, there was no significant increase in the number of MLN CD4 T cells or Tregs (Fig. 1e).

Infection with BRD509-2W1S allows identification of *S.* Tm-specific T cells with 2W1S:I-A^b^ tetramers (Figure S1). In the colon and caecum, 2W1S-specific cells comprise 5-10% of total CD4 T cells (Fig. 1f, Figure S2), but are less frequent in lymphoid tissues including MLNs (Fig. 1g, Figure S2). Tracking 2W1S-specific CD4 T cells over time showed these cells were detectable until at least 90 days p.i. (Fig. 1f-g). The large number of Tconvs specific for 2W1S reflects the immunodominance of the peptide, as previously reported^25–28^. The higher number of 2W1S-specific T cells in the intestinal LP than the MLNs is consistent with the hypothesis that these cells are accumulating at barrier sites to protect against invasion from the intestinal lumen, where the highest bacterial burden is present (Fig. S4a).

### The MLN CD4 T cell response to *Salmonella* is constrained to colon-draining MLNs

The higher bacterial burden and number of 2W1S-specific T cells observed in the large intestine compared to the SI (Fig. 1c, Figure S2b), led to the hypothesis that the MLN CD4 T cell response was focused in the colon and caecum draining MLNs (cMLNs). This hypothesis is consistent with previous work showing that different intestinal sites drain to specific MLNs^29,30^ (Figure S3a). With the aim of improving sensitivity of detecting *S.* Tm responses in the MLNs, a comparison of CD4 T cells in the cMLNs and sMLNs was conducted. Results show that cMLNs but not sMLNs contain an increased number of total cells, CD4 T cells, CXCR3^+^ CD4 T cells and 2W1S-specific T cells following *S.* Tm infection (Figure S3b-c). Together, these data show that the CD4 T cell response to *S.* Tm in the MLN is restricted to the colon and caecum-draining lymph nodes.

### Characterization of 2W1S-specific T cells

To investigate the CD4 T cell response to *S*. Tm infection we initially examined the 2W1S-specific T cell response, and observed that 2W1S-specific T cells homogenously display a Th1 phenotype (Figure S4), consistent with previous reports^28,31,32^. As such, 2W1S-specific cells expressed T-bet but not RORγT or FoxP3, up to and including 60 days p.i. (Fig. S4a-b). Cytokine profiling of 2W1S-specific cells confirms a Th1 phenotype, with cells expressing IFNγ but not IL17A (Fig. S4c-d). It has been previously shown that *S.* Tm-specific CD4 T cells comprise cells that recognize a wide range of peptides, and both *S*. Tm-specific and non-*S.* Tm-specific ‘bystander’ cells play an important role in *Salmonella* clearance and protection from reinfection^31,33–35^. Because of the phenotypic homogeneity of 2W1S-specific cells and the importance of a heterogenous CD4 T cell response to *S*. Tm, we next characterized changes in the total CD4 T cell pool.

### *S*. Tm infection induces an early colonic Th17 response followed by a sustained Th1 response

In contrast to 2W1S-specific cells, total CD4 T cells comprise heterogenous and dynamic populations of Th1 cells, Th17 cells, and Tregs following *S*. Tm infection. To characterize bulk CD4 T cells that includes a mixture of CD4 T cell responding to the bacteria, potential bystander cells, and cells not involved in the response, we analyzed activated conventional T cells (Tconvs) by transcription factor (TF) and cytokine expression. Tconvs were identified as CD4 T cells (gated as described in Fig. 1d) that are CD44^hi^ and FoxP3^**-**^ (Fig. 2a, left). Tconvs were characterized by expression of T-bet, RORγT and GATA3 to identify putative Th1, Th17 and Th2 cells respectively^36^.

**Fig. 2.**
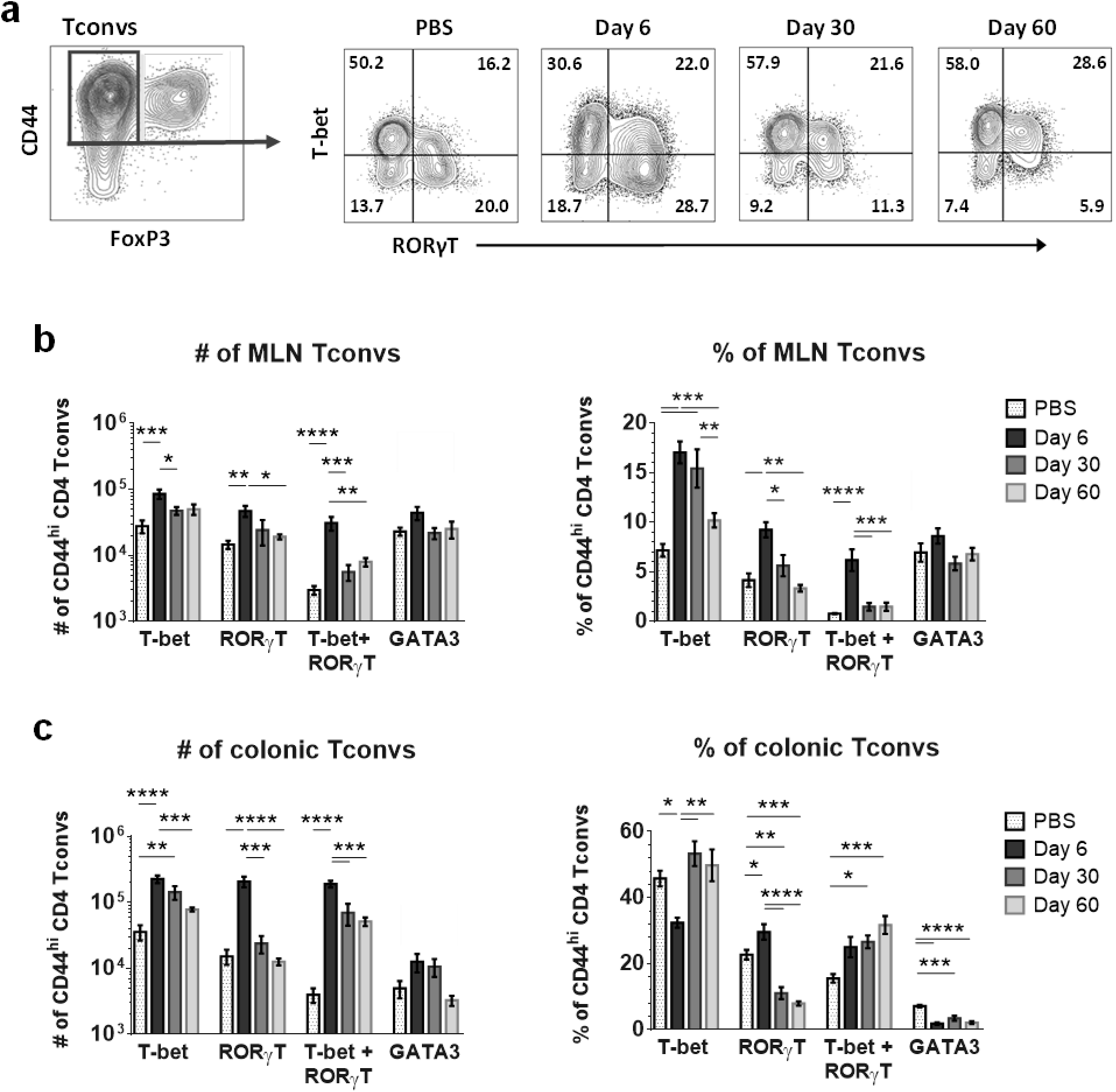
At day 6 p.i. the proportion of Tbet^+^ colonic Tconvs decreases while the proportion of RORyT^+^ Tconvs increases. **a** Representative plots of colonic Tconvs (CD44^hi^ FoxP3^**-**^ CD4^+^ cells, gated left) stained for T-bet and RORγT, following *S.* Tm or mock (PBS) infection. **b** TF expression by cMLN Tconvs, shown as absolute numbers (left) or proportions of Tconvs (right). **c** TF expression by colonic Tconvs, shown as absolute numbers (left) or proportions of Tconvs (right). Means ± SEM are plotted (*n*=3-6 animals/group) and are representative of three independent experiments. Statistical significance calculated by one-way ANOVA with Tukey’s test for each TF. ns, not significant; \**p*<.05; \*\**p*<.01; \*\*\**p*<.001; \*\*\*\**p*<.0001.

The number and proportion of cMLN (Fig. 2b) and colonic LP (Fig. 2c) CD44^hi^ Tconvs are plotted from mock (PBS)-infected mice and *S.* Tm-infected mice examined at 6, 30 or 60 days p.i. In the cMLN, there are increased numbers and proportions of Tconvs that are T-bet^+^, RORγT^+^ and T-bet^+^RORγT^+^ cells at 6 days p.i. (Fig. 2b). Numbers of Tconv subsets return to baseline levels by day 30 p.i., except T-bet^+^ cells, which remain elevated until 60 days p.i. In the colonic LP there is an increased number of Tconvs expressing T-bet, RORγT, or T-bet and RORγT 6 days p.i (Fig. 2c, left). While T-bet^+^, RORγT^+^, and T-bet^+^RORγT^+^ cells increase in number, the increase in RORγT^+^ cells is greater, leading to an increased proportion of RORγT^+^ cells and a decreased proportion of T-bet^+^ and GATA3^+^ cells at day 6 (Fig. 2c, right). This increase in RORγT^+^ Tconvs at day 6 p.i. is transient however, and by day 30 a strong bias towards T-bet^+^ Tconvs is observed. In summary, there is an initial colonic RORγT^+^ response which shifts to a long-lasting T-bet^+^ response. This dynamic CD4 T cell response is observed in the colonic mucosa but not in the cMLN.

### Reciprocal dynamics between colonic Tconvs and Tregs expressing the same TFs

Following characterization of Tconvs by TF, FoxP3^**+**^ Tregs were assessed for expression of Th master TFs T-bet, RORγT and GATA3 (Fig. 3a-b). The number and proportion of colon Tregs expressing these TFs were plotted from mock (PBS)-infected mice or *S.* Tm-infected mice examined at 6, 30 or 60 days p.i. (Fig. 3b). There was an increased number of T-bet^+^ and T-bet^+^RORγT^+^ Tregs at 6 days p.i., but no increase of RORγT^+^ or GATA3^+^ Tregs. Changes in absolute numbers correspond with an increased proportion of T-bet^+^ Tregs and a reduced proportion of RORγT^+^ Tregs at day 6 p.i. While these proportional changes are partially resolved by day 60 p.i., the percentage of Tregs that are T-bet^+^ remain elevated while the proportion of RORγT^+^ Tregs remain lower than mock-infected controls (Fig. 3b, right).

**Fig. 3.**
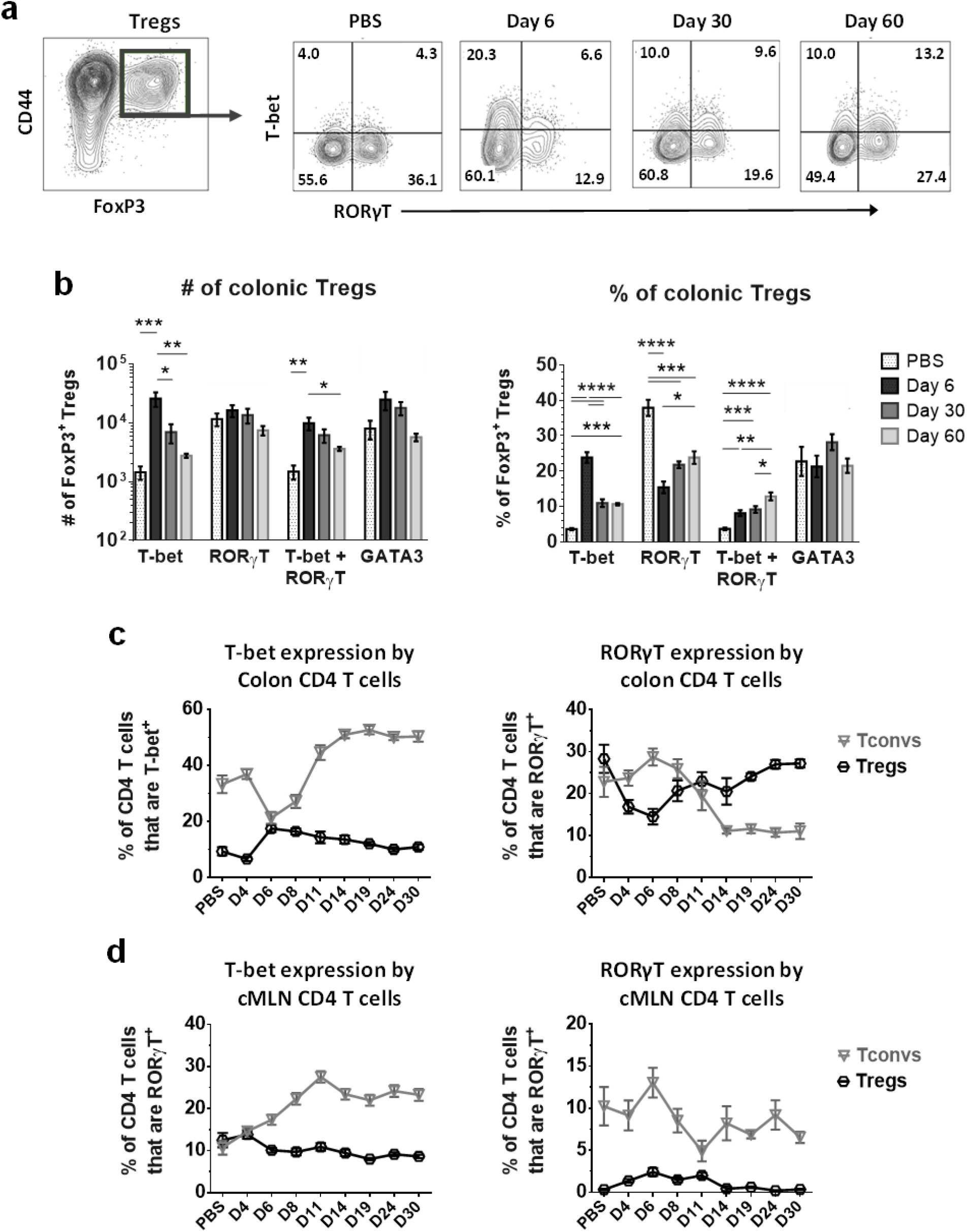
Transcription factor expression by FoxP3^+^ Tregs reveals reciprocity between Tconvs and Tregs expressing T-bet or RORγT in the colon but not the cMLNs. **a** Representative plots of colon Tregs (FoxP3^+^ CD4 T cells, gated left) stained for T-bet and RORγT following mock (PBS) or *S.* Tm infection. **b** Expression of TFs by colonic Tregs are shown as absolute numbers (left) or proportions of Tregs (right). **c** Line charts show changes in in the proportion of colon Tregs and Tconvs expressing T-bet (left) or RORγT (right) during the first 30 days p.i. **d** The expression of T-bet or RORγT by cMLN Tconvs and Tregs are shown. Means ± SEM are plotted (n=3-6 animals/group) and are representative of three independent experiments (**b**). Statistical significance calculated by one-way ANOVA with Tukey’s test for each TF. ns, not significant; \**p*<.05; \*\**p*<.01; \*\*\**p*<.001; \*\*\*\**p*<.0001.

Comparing the proportion of colonic Tconvs and Tregs expressing T-bet or RORγT reveals a reciprocal dynamic between populations expressing the same TF. The early decrease in the proportion of T-bet^+^ Tconvs (Fig. 2c, right) occurs concurrently with an increased proportion of T-bet^+^ Tregs (Fig. 3b, right). Conversely, the transient increase in the proportion of RORγT^+^ Tconvs corresponds with transient decrease in the proportion of RORγT^+^ Tregs.

To further investigate this reciprocity, a more detailed time-course was carried out during the first 30 days p.i. In Fig. 3c-d, the proportions of Tregs that are T-bet^+^ or RORγT^+^ (black lines) are overlaid with the proportion of Tconvs (grey lines) that express the same TFs. These plots highlight a reciprocal dynamic in the colon but not the cMLN. Furthermore, these data show that day 6 p.i. is the peak of the colonic RORγT^+^ Tconv response and that the T-bet^+^ Tconv bias is re-established by day 11 p.i (Fig. 3c). In summary, the early RORγT^+^ Tconv response in the colon coincides with an increased proportion of T-bet^+^ Tregs, and the later T-bet Tconv bias coincides with an increased proportion of RORγT^+^ Tregs.

### Cytokine profiling supports characterization of Th subsets by TF

Next, we assessed IFNγ and IL17A expression to determine whether T-bet^+^ and RORγT^+^ Tconvs are *bona fide* Th1 and Th17 cells, as previously described.^37,38^ At days 6 and 11 p.i., CD4 T cell-generated IFNγ is predominantly expressed by T-bet^+^ Tconvs, and IL17A is predominantly expressed by RORγT^+^ Tconvs (Fig. 4a). The proportion of RORγt^+^ Tconvs cells correlates with the proportion of IL-17A^+^ cells and the proportion of T-bet^+^ Tconvs correlates with IFNγ^+^ cells, in both the colon and MLN (Fig. 4b-c). Furthermore, T-bet^+^ Tconvs predominantly produce IFNγ while RORγT^+^ Tconvs produce IL17A (Fig. 4d). T-bet^+^RORγT^+^ double positive cells are heterogenous in cytokine production and include IFNγ^+^ (Th1), IL17A^+^ (Th17) and IFNγ^+^IL17A^+^ (Th1/17 ‘double positive’) cells. IFNγ/IL17A double positive cells are proportionally the smallest component, comprising <20% of T-bet^+^RORγT^+^ Tconvs in the colon (Fig. 4c). Together, these data support that T-bet and RORγT are appropriate markers for Th1 and Th17 cells in this model.

**Fig. 4.**
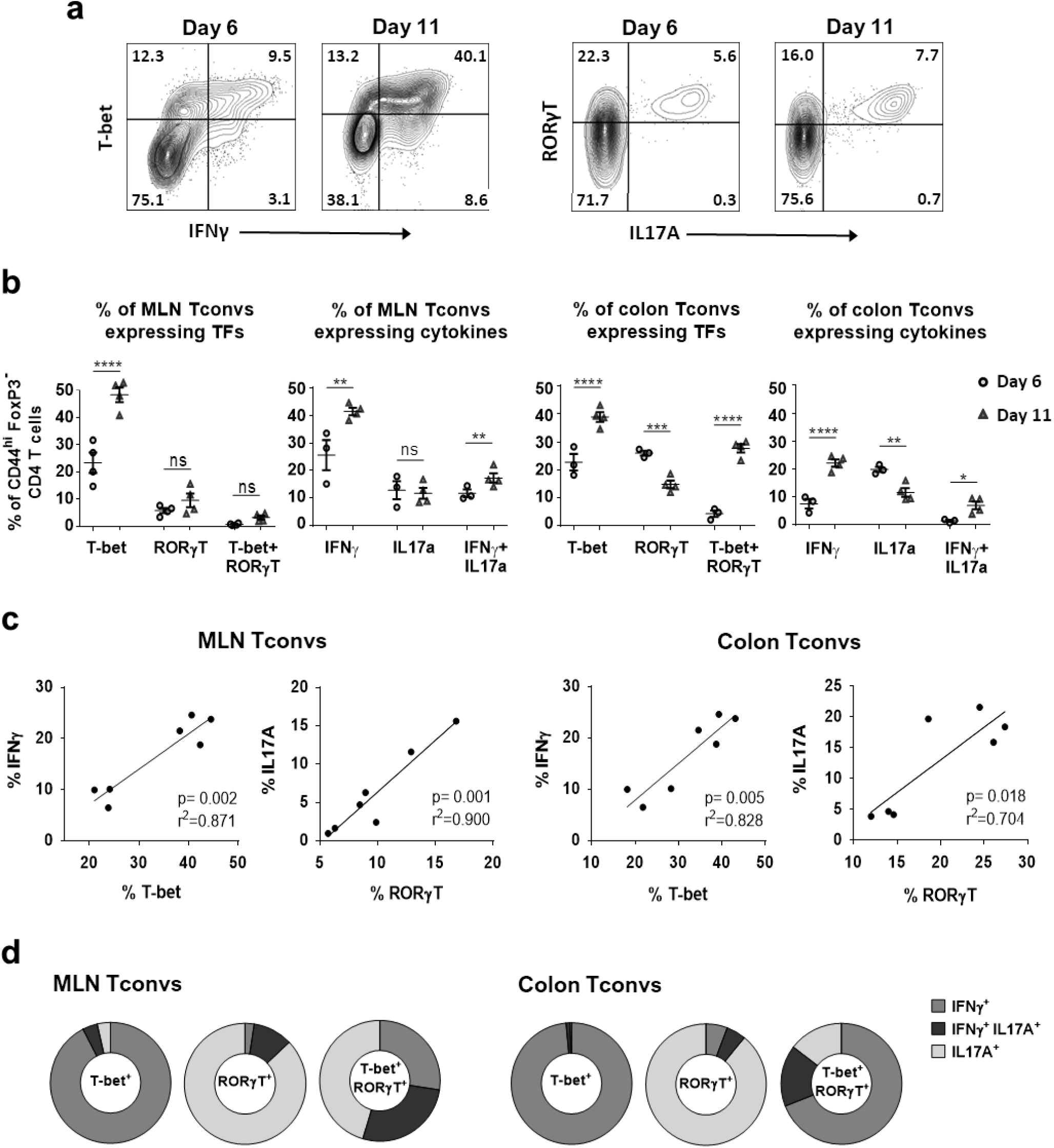
Transcription factor and cytokine expression by Tconvs following *S.* Tm infection. **a** Representative plots of IFNγ and IL-17A expression by *S.* Tm-infected Tconvs at 6 and 11 days p.i. **b** The proportion of MLN and colonic Tconvs that express T-bet and/or RORγT and IFNγ and/or IL-17A are graphed **c** Plots showing the correlation between Tconvs expressing T-bet and IFNγ or RORγT and IL17A. **d** Expression of IFNγ and/or IL17A by Tconvs that are T-bet^+^, RORγT^+^ or T-bet^+^ RORγT^+^ are depicted in donut charts. Means ± SEM are plotted. Data points represent individual animals (n=3-4) from a representative example of two independent experiments. Statistical significance calculated by one-way ANOVA with Holm-Šídák test (b); regression p values calculated by Pearson’s correlation coefficient (c). ns, not significant; *p<.05; **p<.01; ***p<.001; ****p<.0001.

### Effect of bacterial burden on T cell responses following *S.* Tm infection

To understand how the presence of *S.* Tm affects changes in the T cell response, we treated mice with antibiotics to clear the infection. *S.* Tm is known to cause persistent infection in mice and bacterial clearance requires prolonged treatment^28,39^. We employed a previously described enrofloxacin treatment from day 5 to day 29 p.i.^33^. In untreated mice, *S.* Tm CFUs could be recovered in luminal contents and the spleen at day 30 p.i.; following treatment, CFUs could not be recovered from any sites or tissue (Fig. S5a). Treatment reduced the proportion and number of tetramer^+^ T cells at day 30, although the number of 2W1S-specific cells in treated animals remained above levels observed at day 6 (Fig. S5b-c). Enrofloxacin treatment did not change the T-bet^+^ phenotype of 2W1S-specific cells (Fig. S5d), but reduced the proportion of Tbet^+^ Tconv and RORgT^+^ Tregs at day 30 p.i. This indicates that *S.* Tm persistence contributes to the Th1 response, but that the reciprocal dynamic between Tconvs and Tregs is maintained even when bacteria are cleared (Fig. S5e-f).

### Helios^hi^ Tregs can co-express T-bet^+^ and GATA3^+^ but not RORγT

The reciprocal dynamic between Tregs and Tconvs raises questions about the ontogeny and stability of Tregs that express T-bet or RORγT. Helios expression has been proposed to differentiate thymic Tregs (tTregs) from peripherally induced Tregs (pTregs)^40^, but this is controversial as Helios expression has also been reported in pTregs^41–43^. To assess whether Tregs expressing Th TFs co-express Helios, t-SNE plots were generated from colon CD4 T cells 11 days after mock or *S.* Tm infection (Fig. 5). Representative plots of total CD4 T cells (Fig. 5a, left and center) and Tregs (Fig. 5a, right) are shown with gates color-coded to identify corresponding populations. Overlaying FoxP3^+^ Tregs (red), FoxP3^**-**^ CD44^hi^ Tconvs (blue) and FoxP3^**-**^CD44^lo^ naïve T cells (green)(Fig. 5a, left) reveals distinct cluster of Tregs (red), Tconvs (blue) and naïve T cells (green), respectively (Fig. 5b, left). Overlaying T-bet^+^ (light blue), RORγT^+^ (plum), T-bet^+^RORγT^+^ (dark blue) and GATA3^+^ (orange) cells identifies Th1, Th17 and Th2 cells in the Tconv clusters, and T-bet^+^, RORγT^+^ and GATA3^+^ Tregs (Fig. 5b-c, left). Following *S.* Tm infection, the Th1 and T-bet^+^RORγT^+^ population are visibly increased and the Th2 population is almost absent in the Tconv cluster (Fig. 5b-c, left).

**Fig. 5.**
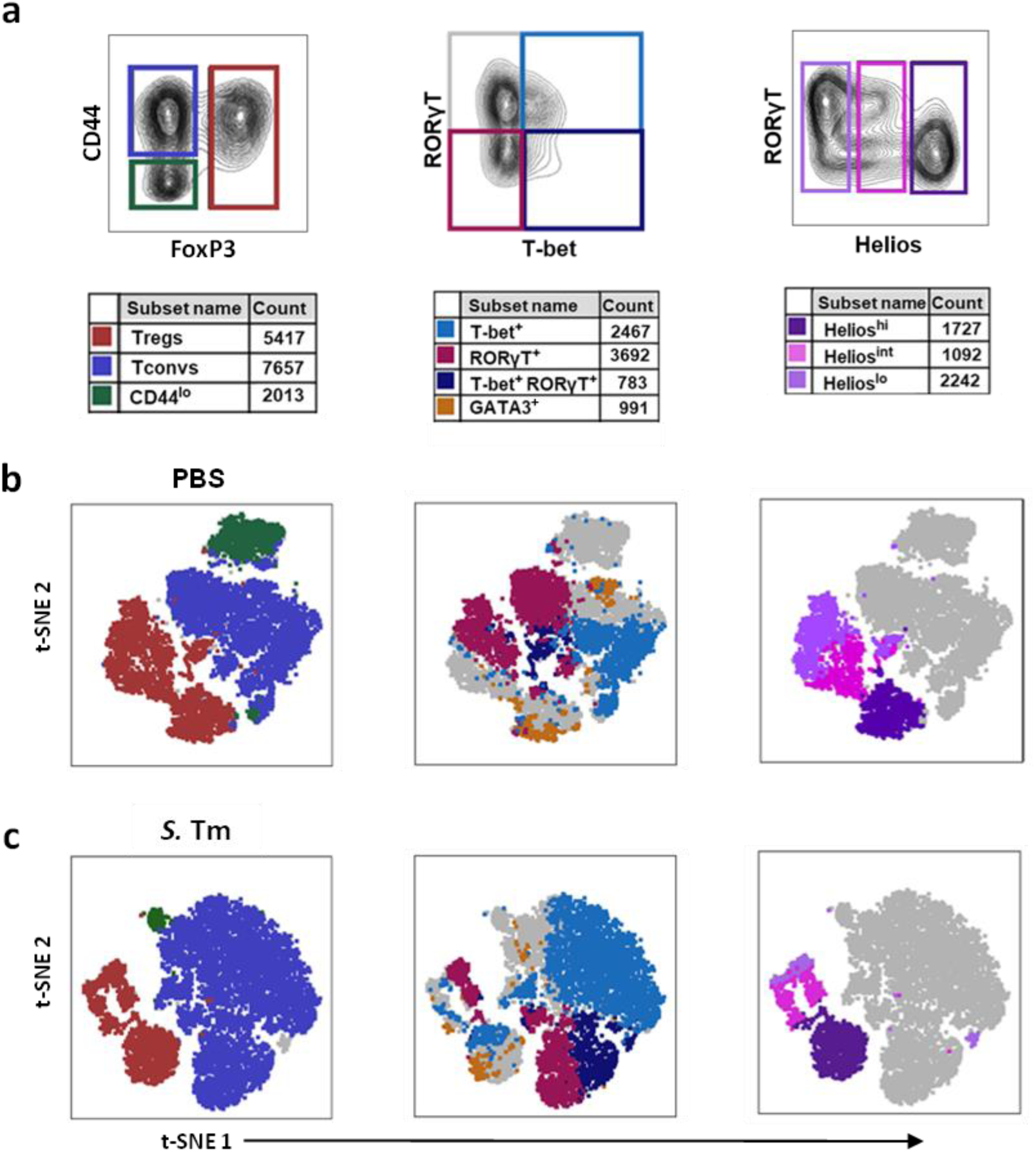
Helios^hi^ Tregs can co-express T-bet or GATA3 but not RORγT. **a** Representative plots show gating and color-coding of colon CD4 T cells that are FoxP3^+^ Tregs, CD44^hi^ Tconvs or FoxP3^**-**^CD44^lo^ naïve T cells (left); T-bet^**+**^, RORγT^**+**^ or T-bet^**+**^ RORγT^**+**^ Tconvs (center); and Helios^**hi**^, Helios^**int**^ or Helios^**lo**^ Tregs (right). Cell counts represent numbers of each population from 2×10^4^ CD4 T cells (gated as described in Fig. 1) and overlaid onto t-SNE plots generated using parameters of CD45, CD3, CD4, CD44, FoxP3, T-bet, RORγT, Helios, GATA3, CCR6 and CXCR3 expression. **b** t-SNE plots are shown for mock (PBS)-infected colons and **c** *S.* Tm-infected colons at day 11 p.i. Grey areas are negative for all markers represented in plots shown above.

Finally, FoxP3^+^ Tregs were gated as Helios^hi^ (dark purple), Helios^int^ (pink) and Helios^lo^ (lilac) populations (Fig. 5a, right). Overlaying these populations reveals that Helios^hi^ Tregs overlap one lobe of the Treg cluster, while Helios^lo/int^ Tregs overlay another (Fig. 5b-c, right). RORγT^+^ Tregs are confined to Helios^lo/int^ cells, GATA3^+^ Tregs are primarily Helios^hi^ cells, and Tbet^+^ Tregs include cells with a range of Helios expression levels. The proportion of Helios^lo^ Tregs is reduced by infection but RORyT^+^ Tregs remain Helios^lo/int^ cells (Fig. 5b-c). Therefore, while *S.* Tm infection alters the proportion of different Treg populations, the co-expression of Th TFs with Helios remains stable.

The non-overlapping expression of Helios and RORγT is consistent with previous work suggesting Helios is a marker of tTregs^40^ and RORγT is a marker of pTregs^22,44^. Defining cell function or ontogeny by marker expression is fraught with difficulties and it is not clear that Helios^hi^ Tregs are *bona fide* tTregs. However, these data in Fig. 5b-c indicate that Tbet^+^ and RORγT^+^ Tregs can be distinguished by Helios expression. In support of this, recent work demonstrating distinct TCR profile between Helios^hi^ and Helios^lo^ Tregs^45^ suggests that T-bet^+^ and RORγT^+^ Tregs are distinct populations instead of different states of differentiation.

### Treg depletion alters the balance of Th1 and Th17 responses following *S.* Tm infection

The reciprocal dynamic between Tregs and Tconvs during *S.* Tm infection (Fig. 3) is consistent with the hypothesis that Tregs expressing Th master TFs selectively target Th subsets and skew the Th response. To determine whether Tregs shape the dynamic Th response to *S.* Tm, Treg depletion experiments were conducted using FoxP3^DTR^ “DEpletion of REGulatory T cell” (DEREG) mice, which allow ablation of FoxP3^+^ Tregs following treatment with diphtheria toxin (DT)^46^. Tregs were efficiently depleted from the peripheral blood and colon after 2 DT treatments on consecutive days. Five days post-treatment there were ∼67% fewer Tregs in the blood, and ∼90% fewer Tregs in the colon than in DT-treated littermate controls (Figure S6a-b). Similar to C57BL/6 mice, Treg-replete DEREG mice exhibit an early Th17 response and later Th1 bias following *S*. Tm infection, and T-bet^+^ and RORgT^+^ Tregs show the same reciprocal dynamic with Tconvs in both strains (Fig. S6d).

In the first depletion experiments, DEREG mice were DT-treated day 1 and 2 p.i., with an endpoint at day 6 p.i. (Fig. 6a). There was no significant difference in weight change between DEREG mice and DTR^-^ (Treg replete) littermates (Figure S7a) nor in *S.* Tm CFUs recovered 6 days p.i. (Figure S7c). The number of total colon cells remained unchanged (Fig. 6b) but both the number of CD4 T cells (Fig. S7d) and CD44^hi^ Tconvs (Fig. 6c) increased following Treg depletion. Although the number of tetramer^+^ cells trended higher, differences were insignificant (Fig. S7e). The increase in CD4 T cells was confined to T-bet^+^ Th1 cells and T-bet^+^RORγT^+^ cells, while the number of RORγT^+^ Th17 cells remained unchanged (Fig. 6d). The increased number of Th1 cells underlies a decreased proportion of Th17 cells and an increased Th1 bias (Fig. 6d, Fig. S7b). The decreased proportion of Th17 cells and increased proportion of Th1 cells following early Treg depletion is robust, also being observed when data from three independent experiments are pooled (Fig. S7b). These pooled data show a colonic Th17 bias in Treg-replete DTR^-^ mice, while there is a Th1 bias in Treg-depleted DEREG littermates (Fig. S7b, right). These data demonstrate that the Th17 bias at 6 days p.i. (Fig. 2b) is dependent on Tregs, which have an increased proportion of T-bet^+^ cells at this timepoint (Fig. 3c, S6d).

**Fig. 6.**
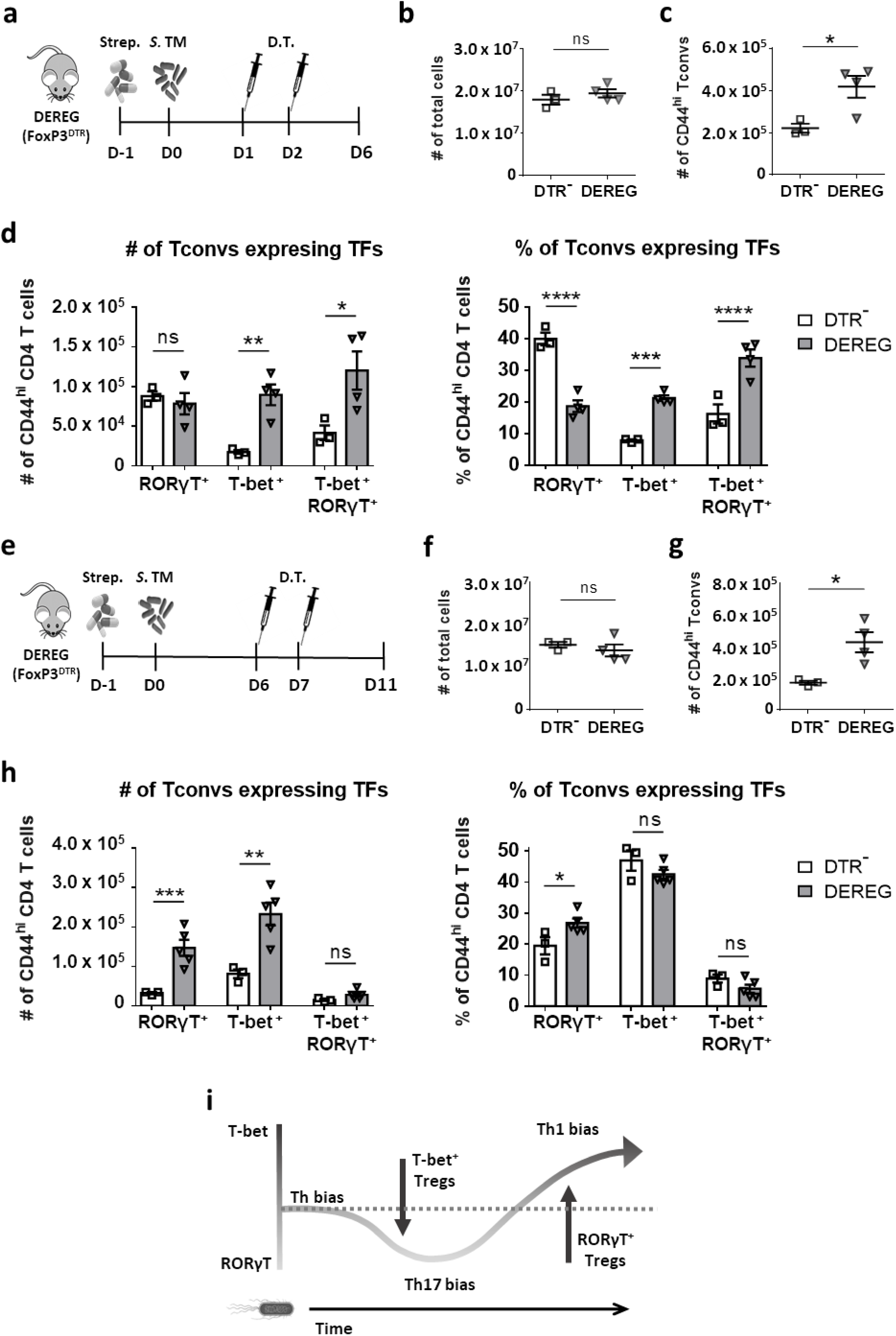
Treg ablation abrogates the early Th17 bias and decreases the later Th1 bias following *S.* Tm infection. **a** Experimental design for Treg depletion of *S.* Tm-infected DEREG mice administered diphtheria toxin (D.T.) at day 1 and 2 p.i., with an endpoint of day 6 p.i. **b** The absolute number of total colonic cells and **c**, CD44^hi^ Tconvs in Treg-depleted DEREG mice and Treg replete DTR^-^ littermates are shown. **d** The absolute number (left) and proportion (right) of colonic Tconvs that are RORγT^+^ or T-bet^+^ are shown at day 6 p.i. **e** Experimental design for Treg depletion of *S.* Tm-infected DEREG mice administered D.T. at day 6 and 7 p.i., with an endpoint at day 11 p.i. **f** The absolute number of total colonic cells and **g**, CD44hi Tconvs in DEREG mice and DTR^-^ littermates are shown. **h** The absolute number (left) and proportion (right) of colon Tconvs that are RORγT^+^ or T-bet^+^ are shown. **i** Schematic of the dynamic Th bias in colonic Tconvs following *S.* Tm infection and the potential for Tregs to shape the CD4 T cell response. The baseline Th bias is indicated by a horizontal dotted line. The potential for Tregs to shape the Th bias is represented by vertical arrows. Data points represent individual animals (n=3-5) from one of three independent experiments. Means ± SEM are plotted. Statistical significance calculated by Mann-Whitney test (**b**,**c**,**f**,**g**) or two-way ANOVA with Holm-Šídák test (**d**,**h**). ns, not significant; *p<.05; **p<.01; ***p<.001; ****p<.0001.

Next, Treg depletion experiments were conducted to determine if Tregs were also required to drive the Th1 bias that becomes prominent by day 11 p.i. (Fig. 2b). DT was administered day 6-7 p.i. and the endpoint was day 11 p.i. (Fig. 6e). As described following earlier DT treatment, later Treg depletion did not cause a significant difference in weight change (Fig. S7f), the number of *S.* Tm CFU (Figure S7h) nor the number of total colon cells (Fig. 6f) between DEREG mice and DTR^-^ littermates. There was an increase in the number of CD4 T cells (Fig. S7i), CD44^hi^ Tconvs (Fig. 6g) and tetramer^+^ 2W1S-specific T cells (Fig. S7j). There was also an increased number of both colonic Th17 and Th1 cells (Fig. 6h, left) but the increase in Th17 cells was greater, reflecting an increased proportion of Th17 cells at day 11 p.i. (Fig. 6h, right). The increased proportion of Th17 cells following Treg depletion is also observed when data from three independent experiments are pooled (Fig. S7g). These pooled data show the Th1 bias in Treg-replete mice at day 11 p.i.is abrogated following Treg depletion (Fig. S7b, right).

To confirm that these changes in Th bias reflect dysregulation of the immune response to *Salmonella*, and not general dysregulation following Treg depletion, we compared the effect of Treg depletion in uninfected DEREG mice (Fig. S6). Four days after DT treatment there was no significant change in Th bias between Treg-depleted DEREG mice and DTR^-^ littermates (Fig. S6e-f). This contrasts with robust changes in Th bias of Treg depleted mice following *S*. Tm infection (Fig. 6d,h; Fig. S7b,g). This indicates Treg depletion does not shift Th bias in the absence of S. Tm infection.

Taken together, these results demonstrate that Tregs are required for both the early Th17 response and a maximal Th1-bias at day 11 p.i. In Fig. 6i, a schematic diagram depicts the Th response following *S.* Tm infection and the effects of Tregs to shape the dynamic Th response.

## DISCUSSION

Here we have developed an *S.* Tm infection model to characterize a dynamic CD4 T cell response. We observe an early colonic Th17 response corresponding with a decreased proportion of RORγT^+^ Tregs and a later, long-lasting Th1 response with a decreased proportion of T-bet^+^ Tregs (Fig. 3b-d) within the total T cells at the infection site. The early Th17 response described here is consistent with work identifying an early expansion and contraction of flagellin-specific Th17 cells in the intestine^33^. The Th1 response to *S.* Tm has been shown to be important for bacterial clearance and protective immunity^5,8,9,47^. The dynamic and site-specific Th response may therefore be optimized to counter a multi-phase infection with changing bacterial targets. However, the factors that control this response are unclear.

The reciprocal dynamic we reveal between Tconvs and Tregs expressing the same TFs is consistent with the hypothesis that Th1 cells are being suppressed by T-bet^+^ Tregs and Th17 cells are being targeted by RORγT^+^ Tregs later after infection. That this dynamic occurs in the colon LP and not the cMLN suggests that reciprocity between Tconvs and Tregs is manifested in the colon itself. Selective suppression of Th1 and Th17 cells by Tregs expressing T-bet or RORγT has been previously reported. Tregs have been shown to upregulate T-bet in response to type 1 inflammation and T-bet^+^ Tregs are required to control Th1-mediated inflammation^24^. In the context of *Listeria* infection, it has been shown that T-bet^+^ Tregs suppress Th1 cells and comprise a stable population that proliferates rapidly during reinfection^21^. It has also been shown that specific intestinal bacteria induce RORγT^+^ Tregs, which limit Th17-mediated colitis, and ablation of Treg-specific STAT3 induces Th17 inflammation^22,23^. Microbiota-specific CD4 T cells have also been shown to be multi-functional and highly plastic^48^. Unlike previous research, here we have characterized a dynamic Th response that is reciprocal to a Treg response. This highlights the potential for Tregs to shape a multi-phase CD4 T cell response in an orchestrated and fine-tuned manner.

To assess the regulation of CD4 T cells, we first assessed changes in *S*. Tm-specific 2W1S:I-A^b^ tetramer^+^ T cells. 2W1S-specific cells maintained a Th1 phenotype in all tissues and timepoints (Fig. S4a-d), in contrast to changes in the bulk CD4 T cell pool. Previous work shows an important role for Tconvs that recognize a wide range of *S*. Tm peptides, as well as non-*S*. Tm-specific non-cognate activated ‘bystander’ cells^31,33–35^, which prompted our focus on the bulk CD4 T cells. As such, tetramer-negative cells are likely a heterogenous population including cells specific for multiple *S*. Tm and non-*S*. Tm antigens. Future work delineating the roles of *S*. Tm-specific and non-*S*. Tm specific cells could help elucidate both the reciprocal dynamics reported here and the functions and interactions of CD4 T cells with different antigen specificity. While the antigen-specificities of tetramer-negative CD4 T cells described here are unknown, we show that *S.* Tm infection drives clear changes in these populations, including increased numbers of Tconvs and Tregs, and changes in their TF expression (Fig. 1d, 2c).

To investigate whether Tregs are responsible for changes in Th bias, Tregs were depleted at critical timepoints. Day 6 p.i. was identified as a peak of the early Th17 response and day 11 p.i. was identified as a timepoint where a Th1 bias was re-established (Fig. 3d). For the day 6 experiment, Tregs were ablated during the period when colonic Tregs are enriched for T-bet^+^ cells; for the day 11 experiment, Tregs were ablated during the period when Tregs comprise a higher proportion of RORγT^+^ cells (Fig. 3b-c, S6c).

At both timepoints Treg depletion increased the number of colonic Tconvs (Fig. 6c,g), consistent with previous reports^49–52^. Treg depletion did not significantly reduce the number of *S.* Tm CFUs recovered (Fig. S7c,h), in contrast to a previous report of reduced bacterial load in DEREG animals systemically-infected with virulent *S.* Tm ^53^. This discrepancy may result from our use of an oral infection model with an attenuated strain, instead of i.v.-infection with a virulent strain. Because the bacterial burden is low in our model (Fig. S5a), ablation of Tregs and shifts in Th bias may not have significant effects on clearance of the few remaining bacteria.

These experiments show that early Treg ablation leads to a selective increase in Th1 cells and prevents the Th17 bias at day 6 p.i. (Fig. 6d, S7b). On the other hand, later Treg ablation leads to an increased number and proportion of Th17 cells and abrogation of the Th1 bias (Fig. 6h, S7g). As such, Tregs, which are enriched for T-bet^+^ cells at early timepoints, are necessary for the early Th17 response. At later timepoints, when Tregs contain an increased proportion of RORγT^+^ cells, Tregs are required for a maximal Th1 bias at day 11 p.i. This indicates that the reciprocal dynamics described here are not just correlative, but that Tregs actively shape the CD4 T cell bias (Fig. 6i). These results are consistent with the hypothesis that T-bet^+^ Tregs selectively suppress the early Th1 response and shift the Tconv response towards a Th17 bias, while RORγT^+^ Tregs inhibit Th17 cells, enhancing the later Th1 response. Future work assessing the effect of selectively depleting Treg subsets will help elucidate the extent to which the dynamic Th response is controlled by selective suppression by Tregs expressing T-bet or RORγT.

The potential for CD4 T cells to up- or down-regulate FoxP3 expression is an alternative mechanism that could underly reciprocity between populations of Tconvs and Tregs expressing the same TFs. Plasticity between regulatory and conventional CD4 T cells has been demonstrated in several contexts. For instance, RORγT^+^ Tregs have been shown to downregulate FoxP3 in the absence of IL-15 and become ‘ex-Tregs’ that drive Th17-mediated colitis ^54^. It has also been shown that Th1 cells have the potential to upregulate FoxP3 to become T-bet^+^ Tregs^55,56^. On the other hand, evidence from lineage tracking models, transfers and *in vitro* experiments suggest that T-bet^+^ Tregs are highly stable and retain FoxP3 expression independent of environmental conditions^21,57^. Because of the stability of T-bet^+^ Tregs, plasticity is a less convincing explanation of reciprocity at later phases of infection. Furthermore, the potential for co-expression of Helios with T-bet but not RORγT by Tregs suggests distinct ontogenies. As such, selective suppression is a possible explanation for both reciprocal dynamics and Treg-dependent Th dynamics, although plasticity between populations of Tconvs or between Tconvs and Tregs may also contribute. Future work using lineage-tracking models and TCR profiling would help delineate the contribution of plasticity between subpopulations.

This research reveals a dynamic Th response in the colon after *S.* Tm infection and a reciprocal dynamic between Tconvs and Tregs expressing the same TFs. Treg depletion experiments show that the early colonic Th17 response and subsequent Th1 bias are dependent on Tregs. In conclusion we highlight that Tregs not only inhibit the Tconv response but shape the CD4 T cell response in a highly nuanced, site-specific and time-dependent manner. These results not only have implications for understanding how the T cell response to *S.* Tm is controlled, but also for how intestinal T cell responses are balanced during infection or inflammation in general.

## MATERIALS AND METHODS

### Mouse strains

Male C57BL/6J mice were purchased from Envigo (Huntingdon, UK) and housed in individually ventilated cages (IVCs) prior to experimental procedures. FoxP3^DTR/eGFP^ DEpletion of REGulatory T cell (DEREG) mice were kindly provided by Prof. Mark Travis (University of Manchester, UK) with permission from Prof. Tim Sparwasser (Medizinische Hochschule Hannover, Germany). DEREG mice were maintained in IVCs. All mice were maintained under specific pathogen-free (SPF) conditions at the University of Glasgow Central Research Facility or Veterinary Research Facility (Glasgow, UK). All procedures were conducted under licenses issued by the UK Home Office under the Animals (Scientific Procedures) Act of 1986 and approved by the University of Glasgow Ethical Review Committee.

### *Salmonella* strains and culture

*S*. Tm BRD509 and BRD509-2W1S were kindly provided by Prof. Stephen McSorley (University of California, Davis, USA). 2W1S strains express the 2W1S (EAWGALANWAVDSA) epitope in frame with OmpC^25,28^. *S.* Tm cultures were streaked out on Luria-Bertani (LB) agar plates before culturing in LB broth for 5 hrs at 37°C in an incubator shaking at 180 rpm. Cultures were then back-diluted 1:10 before static culture at 37°C overnight (O/N). Cultures were then adjusted to an OD_600_ of 1.00, centrifuged at 5,000G for 10 min, and resuspended in sterile phosphate buffered saline (PBS) at an estimated concentration of 1.0-1.5 x 10^9^ CFU/ml. Following infections, actual bacterial dosage was confirmed by plating serial dilutions of *S.* Tm inocula onto LB agar plates.

### *In vivo* infections

6-10 week old mice were pre-treated with 20 mg streptomycin (Sigma-Aldrich, USA) suspended in 10 µl sterile PBS by oral gavage 24 hrs before *S.* Tm infection. Mice were infected by administration of 100 µl of *S.* Tm inoculum or PBS by oral gavage. For infection time-course experiments (Fig. 1d-f, 2b-c, 3b-d), control animals were PBS (mock)-infected at the same time as *S.* Tm-infected animals. At each timepoint, *S*. Tm- and mock (PBS)-infected animals were analyzed. Mock infected controls from each timepoint are combined into single control groups shown in time-course graphs.

### Bacterial recovery

Single cell suspensions from tissues were pelleted by centrifuging at 400 G for 5 mins and resuspended in 0.1% Triton X-100 (Sigma-Aldrich) in PBS and incubated at room temperature (RT) for 10 mins. Samples were then washed and pelleted before being resuspended in PBS, serially diluted and plated on MacConkey agar No. 2 (ThermoFisher, UK) containing 5 µg/ml streptomycin (Sigma-Aldrich) and incubated O/N at 37°C before CFUs were calculated. Bacteria were recovered from feces and caecal contents, which were collected, aliquoted into 100 µg samples, homogenized and serially diluted and plated on MacConkey agar plates as described above.

### Enrofloxacin treatment

Antibiotic treatment of *S.* Tm-infected mice was carried out by adding enrofloxacin (Bayer, Germany) to drinking water at 2 mg/ml. Enrofloxacin treatment was provided from day 5-29 post infection (p.i.) and water was replaced every 72 hrs.

### Tissue harvest and processing

External fat, Peyer’ patches (PP) and caecal patches (CP) were removed from intestinal samples and the remaining tissue was chopped and washed in HBSS with 2 mM EDTA (Gibco, UK). Samples were then incubated at 37°C shaking at 205 rpm for 10 mins, washed in EDTA buffer and the process was repeated twice. EDTA incubations and washes were repeated thrice before digestion. Digest enzyme cocktails were prepared in complete RPMI media (RPMI 1640 with 100 µg/ml streptomycin, 100 U/m penicillin, 2 mM L-Glutamine and 50 µm 2-Mercaptoethanol) with 10% FCS (all Gibco). Colon and caecal tissue were digested in an enzyme cocktail of 0.45 mg/ml collagenase V (Sigma-Aldrich), 0.65 mg/ml collagenase D (Roche, Switzerland), 1.0 mg/ml dispase (Gibco) and 30 µg/ml DNAse (Roche). Small intestines were digested with 0.5 mg/ml collagenase V (Sigma-Aldrich). Tissues were incubated at 37°C in an incubator shaking at 205 rpm for 15-20 mins. Following digests, samples were filtered through 100 µm filters, washed twice with buffer (PBS with 2% FCS and 2 mm EDTA) and filtered through a 40 µm filter.

Lymph nodes, Peyer’s patches (PPs) and caecal patches (CPs) were washed in HBSS and chopped into ∼1 mm pieces, passed through a 40 µm filter and suspended in buffer. Spleens were cleaned and fat was removed before passing through a 40 µm filter with FACS buffer. Samples were suspended in Ammonium-Chloride-Potassium (ACK) lysing buffer (ThermoFisher) for 3-5 mins on ice, before washing and resuspending in FACS buffer.

### Staining for flow cytometry

2W1S:I-A^b^ tetramer was kindly provided by the NIH tetramer core facility (Atlanta, GA). Following infection with *S.* Tm-2W1S strains, tissues were processed as described above. Samples were transferred into 96 well round bottom plates and resuspended in 25µl of tetramer mix containing complete RPMI with 10% FCS, 7 µg/ml 2W1S:IA^b^ tetramer, 20 µl/ml mouse serum and 10 µl/ml Fc block (CD16/32) (Biolegend, CA). Plates were incubated for 2 hrs at 37°C. Following tetramer staining, samples were stained with fixable viability dye at a 1 µl/ml in PBS. Next, antibodies for extracellular markers were prepared at 1:100 or 1:200 dilutions in FACS buffer, as specified in the list of antibodies used (Figure S7). Surface staining was performed at 4°C for 30 mins before washing and resuspension.

Transcription factor (TF) staining was carried out after cells were fixed using the eBioscience FoxP3 transcription factor staining kit (ThermoFisher) or BD Cytofix (BD Biosciences, Belgium) according to manufacturer’s instructions. Cells were fixed for 1 hr at room temperature (RT) in the dark before washing and resuspending in eBioscience FoxP3 permeabilization buffer (PB) (ThermoFisher). Following fixation, TF antibodies were diluted in PB at concentrations listed in Figure S7, added to samples and incubated O/N at 4°C.

For cytokine staining, cells were treated with eBioscience cell stimulation cocktail (ThermoFisher), containing phorbol 12-myristate 13-acetate (PMA), ionomycin, brefeldin A and monensin, according to manufacturer instruction. Samples were then surface stained and fixed with BD Cytofix (BD Biosciences) according to manufacturer’s instructions and stained with cytokine antibodies diluted for 2 hrs at RT. Following cytokine staining, cells were washed and resuspended in FACS buffer.

### Flow cytometry

Samples were analyzed using a BD LSR Fortessa analyzer or sorted using a BD FACSAria IIu or III (BD Biosciences) at the Institute of Infection, Immunity and Inflammation Flow Core Facility (University of Glasgow).

### Regulatory T cell depletion

Diphtheria toxin (DT) (Sigma-Aldrich) was suspended in PBS at a concentration of 10 µg/ml. DT was administered intraperitoneally (IP) twice, at a 24 hr interval, at a concentration equivalent to 30 ng/g mouse weight. DT was administered to DTR^**+**^ DEREG mice and DTR^−^ littermates as Treg-replete controls.

## ACKNOWLEDGEMENTS

Thanks to Prof. Stephen McSorley for *S.* Tm strain BRD509-2W1S, Profs. Mark Travis and Tim Sparwasser for DEREG mice, and the National Institute of Health Tetramer Core for 2W1S:IA^b^ tetramer. Thanks also to Diane Vaughan and the Institute of Infection, Immunity and Inflammation Flow Core Facility. Kind thanks to Drs Claire McIntyre, Josh Gray and Shaima Al-Khabouri for assistance with experiments. Funding was provided by the Medical Research Council (MRC/1654776 and MR/N023625/1).

## AUTHOR CONTRIBUTIONS

S.L.C. Conceived of study, designed experiments, conducted experiments, analysed data and prepared manuscript. S.W.F.M. and M.K.L.M Conceived of study, designed experiments, analysed data and edited manuscript. A.B.B. Conducted experiments. D.M.W. contributed to experiments involving *S.* Tm infections and edited the final manuscript.

## Notes

**CONFLICT OF INTEREST** None declared.

### Competing Interest Statement

The authors have declared no competing interest.

