## Supplemental Figures for "Regulatory T cells control the dynamic and site-specific polarization of total CD4 T cells following *Salmonella* infection" (from another version of this preprint: version 2, 2020-04-09)

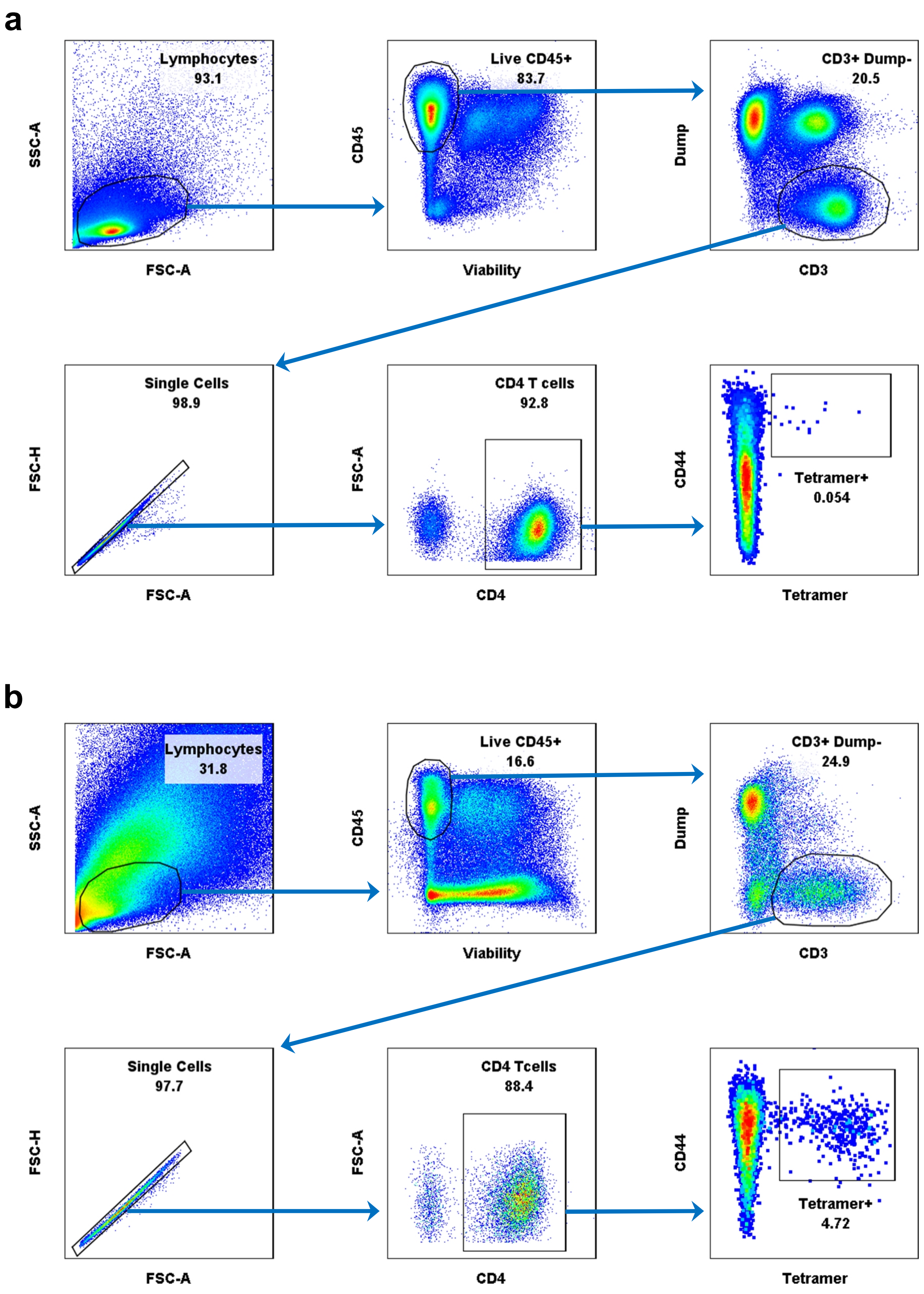


**Figure S1** Gating strategy for 2W1S:IA$^{b}$ tetramer$^{+}$ CD4 T cells in the MLNs and colon. Representative plots from BRD509-2W1S-infected mice 6 days p.i. Tetramer$^{+}$ cells are identified from single cell suspension of **a**, MLN and **b**, colonic cells. Cells are identified as live, single CD45$^{+}$, CD3$^{+}$, Dump (MHCII, B220, CD8, CD64)$^{-}$, CD4$^{+}$, CD44$^{hi}$ and tetramer$^{+}$ cells.

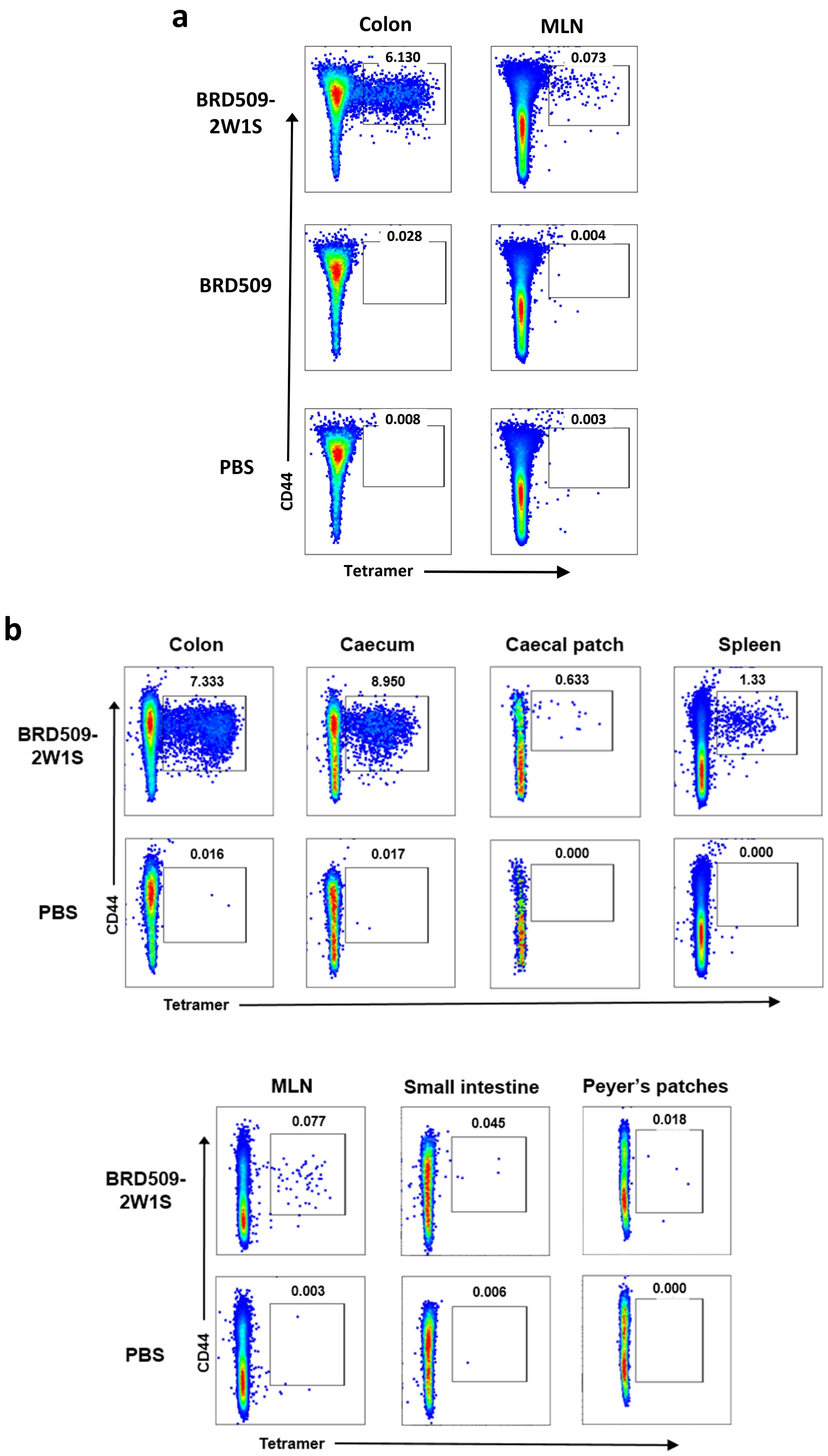


**Figure S2** 2W1S-specific CD4 T cells are detected in multiple intestinal and lymphoid sites. **a** Representative plots of 2W1S:IA$^b$ tetramer$^+$ CD4 T cells from the colon and MLN of animals infected with *S.* Tm strain BRD509-2W1S, the non-2W1S-expressing BRD509 parental strain, or mock infected with PBS. **b** Representative plots of tetramer$^+$ cells from 7 tissues at day 30 post-infection with BRD509-2W1S (top) or mock-infected with PBS (bottom).

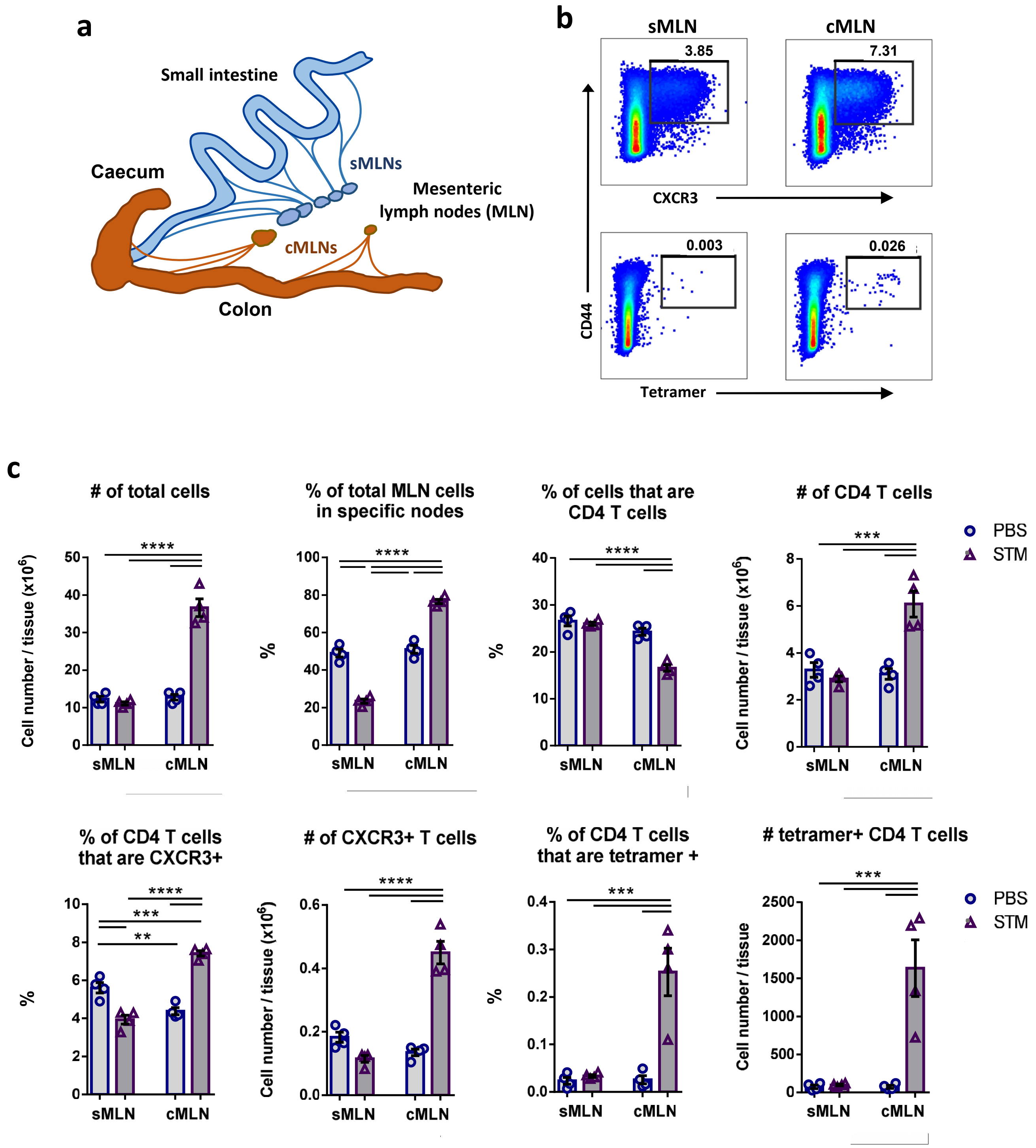


**Figure S3** The MLN CD4 T cell response 6 days p.i. is confined to colon and caecum draining nodes. **a** Schematic representation of intestines and MLN, including small intestine draining MLNs (sMLN), and colonic draining MLNs (cMLN). **b** Representative plots of CD4 T cells from sMLN (left) and cMLN (right) that are CXCR3$^+$ (top) and tetramer$^+$ (bottom), 6 days p.i. **c** Graphs showing the number and proportion of total cells, CD4 T cells, and CD4 T cells that are CXCR3$^+$ or tetramer$^+$ at day 6 p.i. Data points represent individual animals (n=4). Mean ± SEM are plotted. Statistical significance calculated by a one-way ANOVA with Tukey's test. *p<.05; **p<.01; ***p<.001; ****p<.0001.

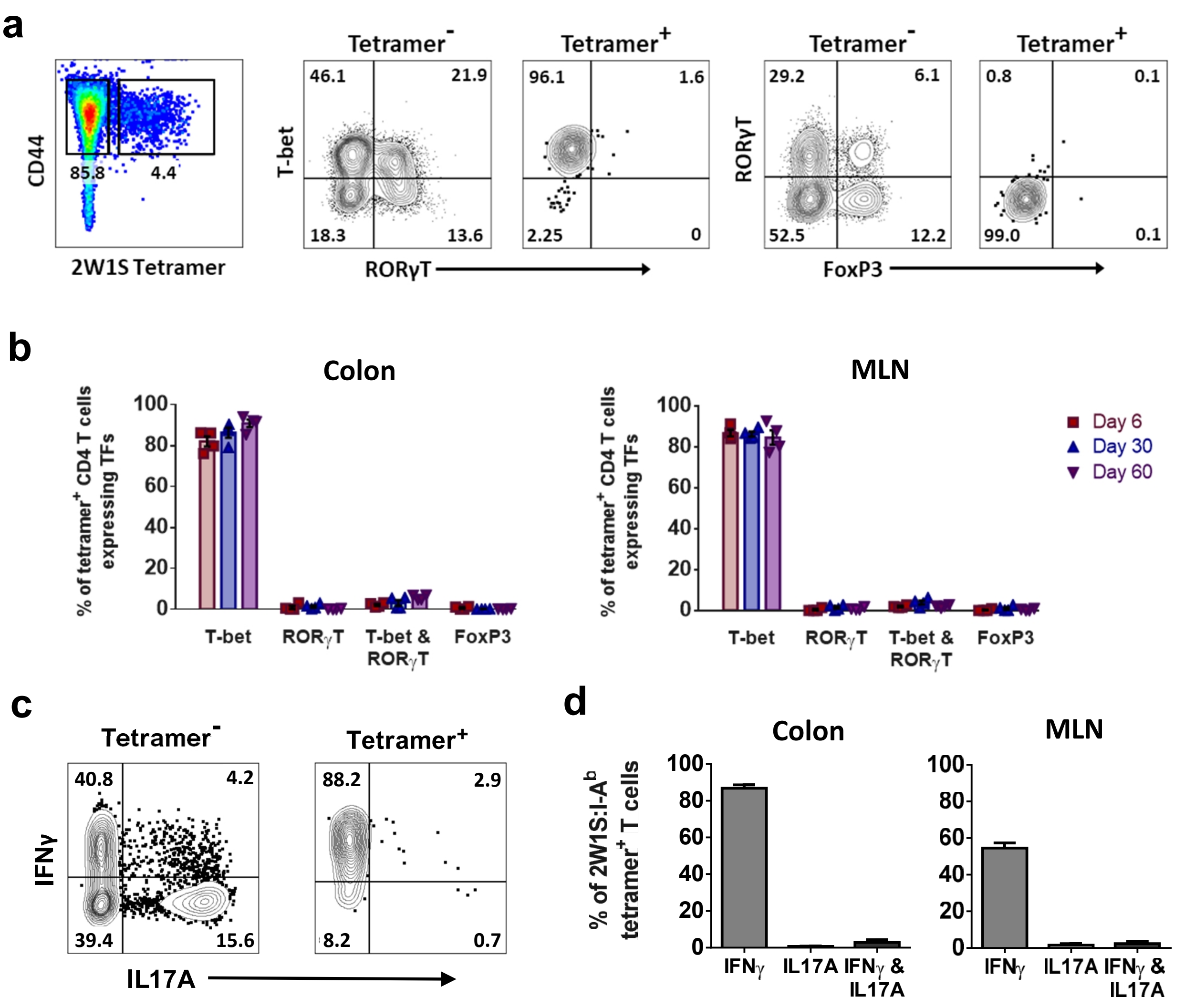


**Figure S4** 2W1S:I-Ab tetramer⁺ CD4 T cells in the colon and MLN express T-bet and IFNγ but not RORγT, FoxP3 or IL17A. **a** Representative plots of TF expression by colonic CD4 T cells that are tetramer⁻ or tetramer⁺. **b** Tetramer⁺ cells in the colon (left) and MLN (right) are predominantly T-bet⁺ and do not express RORγT or FoxP3 at day 6, day 30 and day 60 p.i. **c** Representative plots of IFNγ and IL-17A expression by 2W1S-specific CD4 T cells 11 days p.i. **d** The proportion of 2W1S:I-Ab tetramer⁺ CD4 T cells expressing IFNγ and/or IL17A in the colon (left) and MLN (right) are shown 11 days p.i. Means ± SEM are plotted.

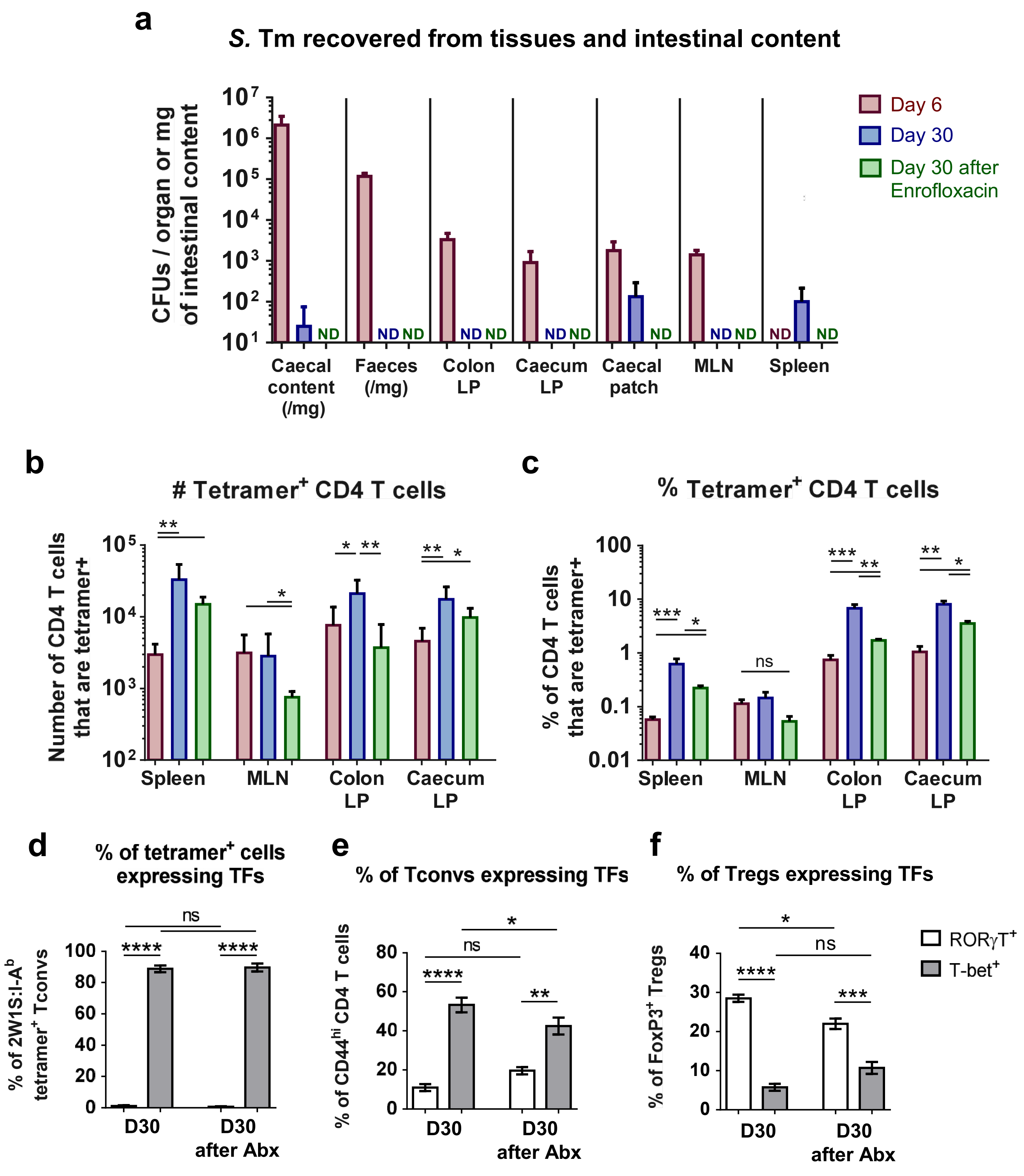


**Figure S5.** Enrofloxacin treatment for 3 weeks eliminates any detectable *Salmonella* CFUs, reduces the number of tetramer$^+$ CD4 T cells, and partially reduces the Th1 bias at day 30 p.i. **a** Samples of caecal content, faeces and tissues were plated following harvest and the number of *S.* Tm CFUs recovered / organ or mg of intestinal content was plotted. **b** The proportion and **c** absolute number of CD4 T cells that are tetramer$^+$ at day 6, 30 or 60 p.i. is shown following 24 days of treatment with enrofloxacin in drinking water from 6 days p.i. The proportion of 2W1S-specific T cells (**d**), Tconvs (**e**) and Tregs (**f**) expressing RORγT or T-bet are shown from enrofloxacin-treated or untreated mice at day 30 p.i. Data is from one experiment (n=3-6). Mean ± SEM are plotted. Statistical significance calculated for each tissue by a one-way ANOVA with Tukey's test. ns, not significant; *p<.05; **p<.01; ***p<.001; ****p<.0001; Abx, antibiotics; ND, not detected).

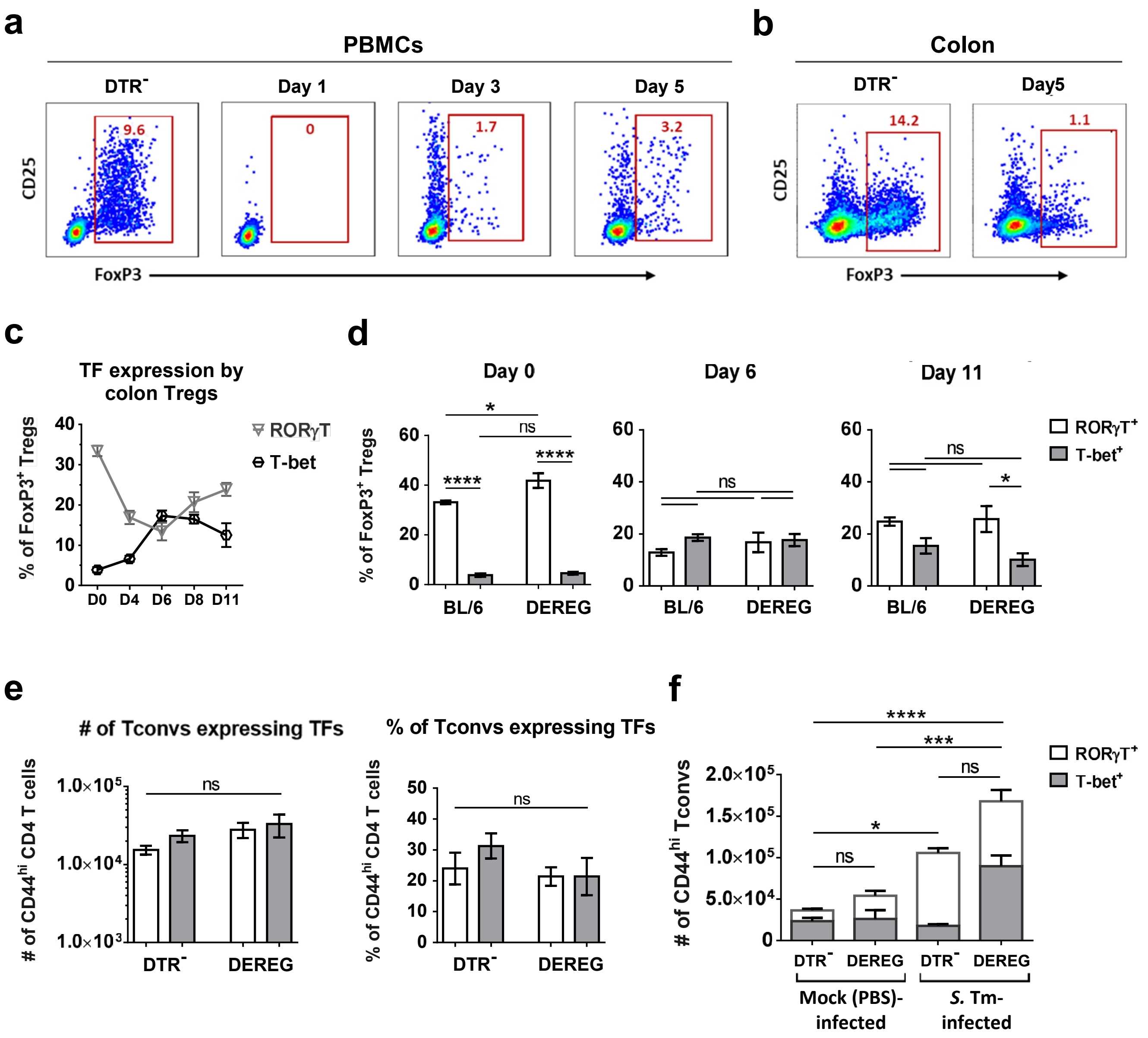


**Figure S6** Diphtheria toxin (D.T.) treatment of DEREG mice depletes Tregs in PBMC and colonic lamina propria; Tregs in untreated DEREG mice express similar levels of T-bet and RORγT as C57BL/6 mice; and Treg ablation in mock-infected mice does not significantly shift the Th1-Th17 bias nor increase colonic Tconv numbers, as observed following *S.* Tm infection. **a** Representative plots of CD4 T cells from PBMCs are shown from a WT littermate (left) and DEREG mouse at 1, 3 or 5 days following 2 D.T. treatments, 24 hrs apart. **b** Representative plots show the proportion of colon CD4 T cells that are FoxP3$^{+}$ 5 days after D.T. treatment. CD4 T cells are identified as described in Fig. 1d. Gate numbers represent the proportion of CD4 T cells that are FoxP3$^{+}$. **c** The proportion of colon Tregs from C57BL/6 mice that express T-bet or RORγT are plotted following *S.* Tm infection. **d** Graphs compare Treg expression of T-bet or RORγT between C57BL/6 mice and Treg replete (untreated) DEREG mice at day 0, 6 and 11 following *S.* Tm infection. **e** The absolute number (left) and proportion (right) of colonic CD44$^{hi}$ Tconvs that are RORγT$^{+}$ or T-bet$^{+}$ are shown 4 days following D.T. treatment in uninfected DEREG mice or wild type (WT) littermates. **f** The number of RORγT$^{+}$ and T-bet$^{+}$ Tconvs are stacked to show changes in cell numbers 4 days following Treg depletion in mock- or *S.* Tm-infected mice 6 days p.i. Data represent n=3-8/group. Means ± SEM are plotted. Statistical significance calculated by one-way ANOVA (**d**, **e**) or two-way ANOVA with Holm-Šídák test of compiled columns (**f**) . ns, not significant; *p<.05; **p<.01; ***p<.001; ****p<.0001.

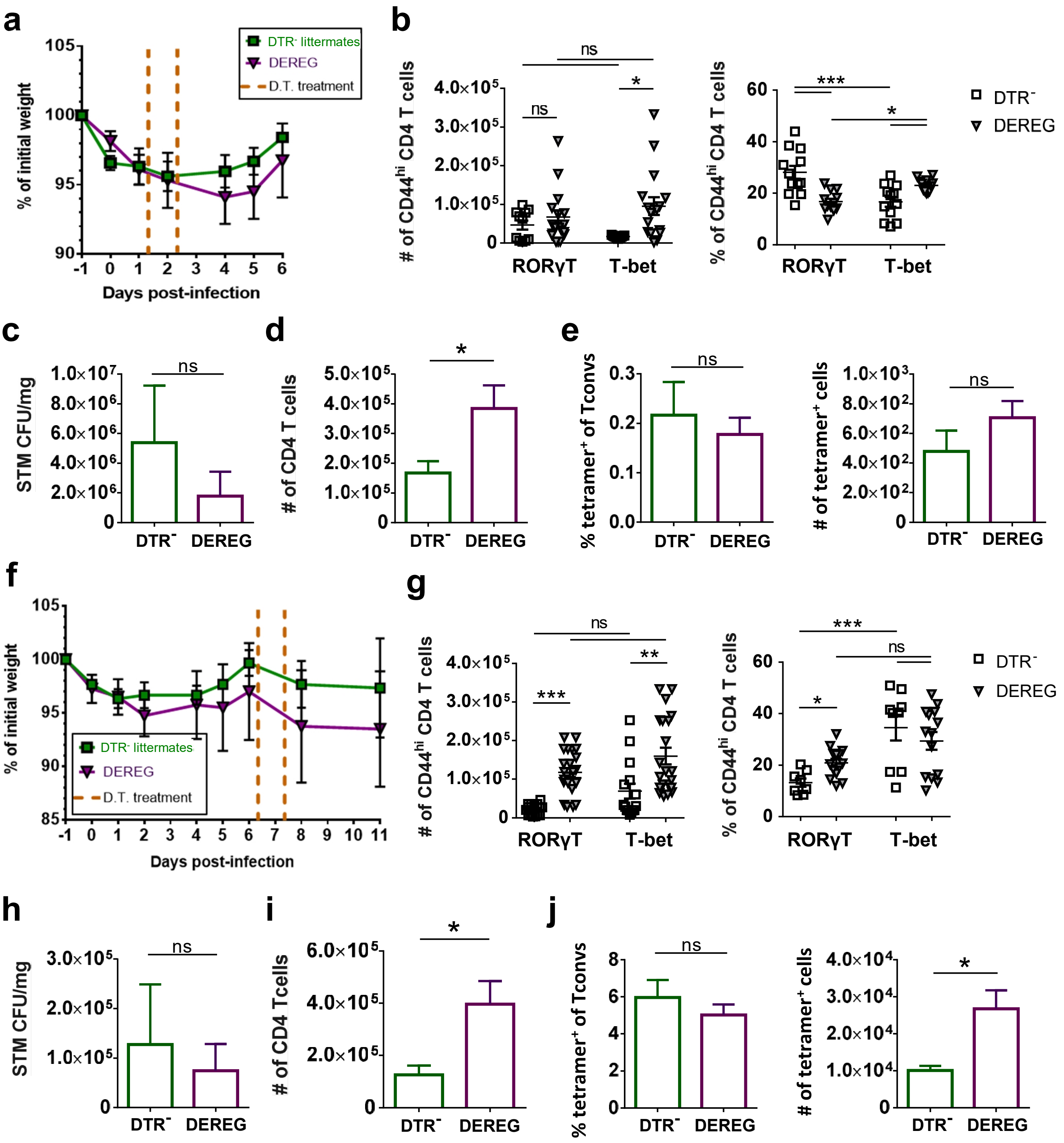


**Figure S7** Treg depletion of *S.* Tm-infected mice does not cause significant weight loss, change faecal *S.* Tm burden or tetramer$^+$ cells, but consistently shifts Th bias. **a** Weight loss in DEREG mice and WT littermates following *S.* Tm infection and D.T. treatment (represented by dashed lines) at day 1-2 p.i. **b** Pooled data showing the number (left) and proportion (right) of CD44$^{hi}$ CD4 Tconvs expressing RORγT or T-bet at day 6 p.i. **c** STM CFU recovered from faeces at day 6 p.i. **d** The number of colonic CD4 T cells in DEREG mice and WT littermates are shown. **e** The proportion (left) and number (right) of colonic Tconvs that are tetramer$^+$ are shown. **f** Weight loss shown following STM infection and D.T. treatments at day 6-7 p.i. **g** Pooled data shows the number (left) and proportion (right) of Tconvs expressing RORγT or T-bet at day 11 p.i. **h** *S.* Tm CFU recovered from faeces at day 11 p.i. **i** The number of CD4 T cells, **j** The proportion (left) and number (right) of colonic Tconvs that are tetramer$^+$ are shown. Data are representative examples of three independent experiments (n=3-5) (**c**-**e**, **h**-**j**) or are pooled from 3 independent experiments (**b, g**). Means ± SEM are plotted. Statistical significance calculated by Mann-Whitney test (**c**-**e**, **h**-**j**) or one-way ANOVA with Holm-Šídák test (**b**, **g**). ns, not significant; *p<.05; **p<.01; ***p<.001.

**Figure S8 Antibodies used for flow cytometry**

| Marker | Fluorochrome | Clone | Concentration | Manufacturer |
|---|---|---|---|---|
| 7AAD | N/A | N/A | 5µl/sample | Biolegend |
| B220 | BV510 | RA3-6B2 | 1:200 | Biolegend |
| CCR6 | BV605 | 29-2L17 | 1:100 | Biolegend |
| CD25 | PE/Cy7 | PC61 | 1:200 | Biolegend |
| CD3 | BV605 | 17A2 | 1:100 | Biolegend |
| CD3ε | AF700 | eBio500A2 | 1:100 | eBioscience |
| CD3ε | BUV395 | 145-2C11 | 1:100 | BD Biosciences |
| CD4 | BUV805 | GK1.5 | 1:200 | BD Biosciences |
| CD4 | AF647 | GK1.5 | 1:200 | Biolegend |
| CD4 | PE/Cy7 | GK1.5 | 1:200 | Biolegend |
| CD44 | BV605 | IM7 | 1:100 | Biolegend |
| CD44 | BV785 | IM7 | 1:100 | Biolegend |
| CD45 | BV785 | 30-F11 | 1:200 | Biolegend |
| CD45 | AF700 | 30-F11 | 1:200 | Biolegend |
| CD8α | BV510 | 53-6.7 | 1:200 | Biolegend |
| CXCR3 | BV421 | CXCR3-173 | 1:100 | Biolegend |
| CXCR3 | BV605 | CXCR3-173 | 1:100 | Biolegend |
| F4/80 | BV510 | BM8 | 1:200 | Biolegend |
| Fixable viability | eFluor 780 | N/A | 1:1000 | Invitrogen |
| FoxP3 | APC | FJK-16s | 1:100 | eBioscience |
| FoxP3 | PE | FJK-16s | 1:100 | eBioscience |
| FoxP3 | AF700 | FJK-16s | 1:100 | Invitrogen |
| GATA3 | PerCP/Cy5.5 | TWAJ | 1:100 | eBioscience |
| Helios | AF488 | 22F6 | 1:100 | BD Biosciences |
| Helios | PerCP/Cy5.5 | 22F6 | 1:100 | Biolegend |
| I-A/I-E | BV510 | M5/114.15.2 | 1:200 | Biolegend |
| IFNγ | PE | 3E4 | 1:100 | eBioscience |
| IFNγ | PerCP/Cy5.5 | XMG1.2 | 1:100 | Biolegend |
| IL-17A | BV605 | TC11-18H10.1 | 1:100 | Biolegend |
| RORγT | PE | AFKJS-9 | 1:100 | eBioscience |
| RORγT | APC | B2D | 1:100 | Invitrogen |
| T-bet | PE/Cy7 | eBio4B10 | 1:100 | eBioscience |
